# Spatially bivariate EEG-neurofeedback can manipulate interhemispheric rebalancing of M1 excitability

**DOI:** 10.1101/2021.12.15.472763

**Authors:** Masaaki Hayashi, Kohei Okuyama, Nobuaki Mizuguchi, Ryotaro Hirose, Taisuke Okamoto, Michiyuki Kawakami, Junichi Ushiba

## Abstract

Human behavior requires interregional crosstalk to employ the sensorimotor processes in the brain. Although some external neuromodulation tools have been used to manipulate interhemispheric sensorimotor activity, a central controversy concerns whether this activity can be volitionally controlled. Experimental tools lack the power to up- or down-regulate the state of the targeted hemisphere over a large dynamic range and, therefore, cannot evaluate the possible volitional control of the activity. We overcame this difficulty by using the recently developed method of spatially bivariate electroencephalography (EEG)-neurofeedback to systematically enable participants to manipulate their bilateral sensorimotor activities. Herein, we report that bi-directional changes in ipsilateral excitability to the imagined hand (conditioning hemisphere) affect interhemispheric inhibition (IHI) assessed by paired-pulse transcranial magnetic stimulation paradigm. In addition, participants were able to robustly manipulate the IHI magnitudes. Further physiological analyses revealed that the self-manipulation of IHI magnitude reflected interhemispheric connectivity in EEG and TMS, which was accompanied by intrinsic bilateral cortical oscillatory activities. Our results provide clear neuroscientific evidence regarding the inhibitory interhemispheric sensorimotor activity and IHI manipulator, thereby challenging the current theoretical concept of recovery of motor function for neurorehabilitation.

## Introduction

Projection neurons wire the brain over long distances and provide a network between different brain regions. In particular, both hemispheres are structurally connected by transcallosal projections and exhibit functional cross talks (Hofer and Frahm, 2006; Meyer et al., 1995); such interhemispheric interaction is essential for higher order cognitive and sensorimotor brain functions. An early argument for interhemispheric interaction, rather than the specific processes performed by each brain area, was that dynamic interplay via the callosum not only allows for simple coordination of processing between the hemispheres but also has profound effects on attentional functioning (Banich, 1998). In the motor domain, neurons in the monkey primary motor cortex are fired bilaterally and motor signals from the two hemispheres interact during unimanual motor tasks (Ames and Churchland, 2019); in the human motor cortex, measuring these neural activities via electroencephalography (EEG) and paired-pulse transcranial magnetic stimulation (TMS) recently revealed that the two hemispheres act together and the related cortical oscillatory activity influences the inhibitory interhemispheric brain network (Picazio et al., 2014; Stefanou et al., 2018).

A critical issue is whether a specific interhemispheric activity can be volitionally controlled. Previous studies have focused on inhibitory interhemispheric sensorimotor network, represented by interhemispheric inhibition (IHI), from movement-related manner (Duque et al., 2007, 2005; Murase et al., 2004), and passive neuromodulation effects by externally administered intervention using transcranial direct current stimulation (tDCS) or repetitive transcranial magnetic stimulation (rTMS) (Boddington and Reynolds, 2017; Gilio et al., 2003; Williams et al., 2010). Due to the experimental limitations related to these observational or open-loop paradigms, much less is known about the effects of changes in sensorimotor activity patterns in both hemispheres on IHI, and the possibility of consciously self-manipulating the inhibitory interhemispheric sensorimotor activity. Therefore, the identification of bilateral sensorimotor activity patterns that underlie IHI and the potent self-manipulator of IHI should be of great value for the understanding of human sensorimotor neural plasticity (Figure 1A).

**Figure 1.**
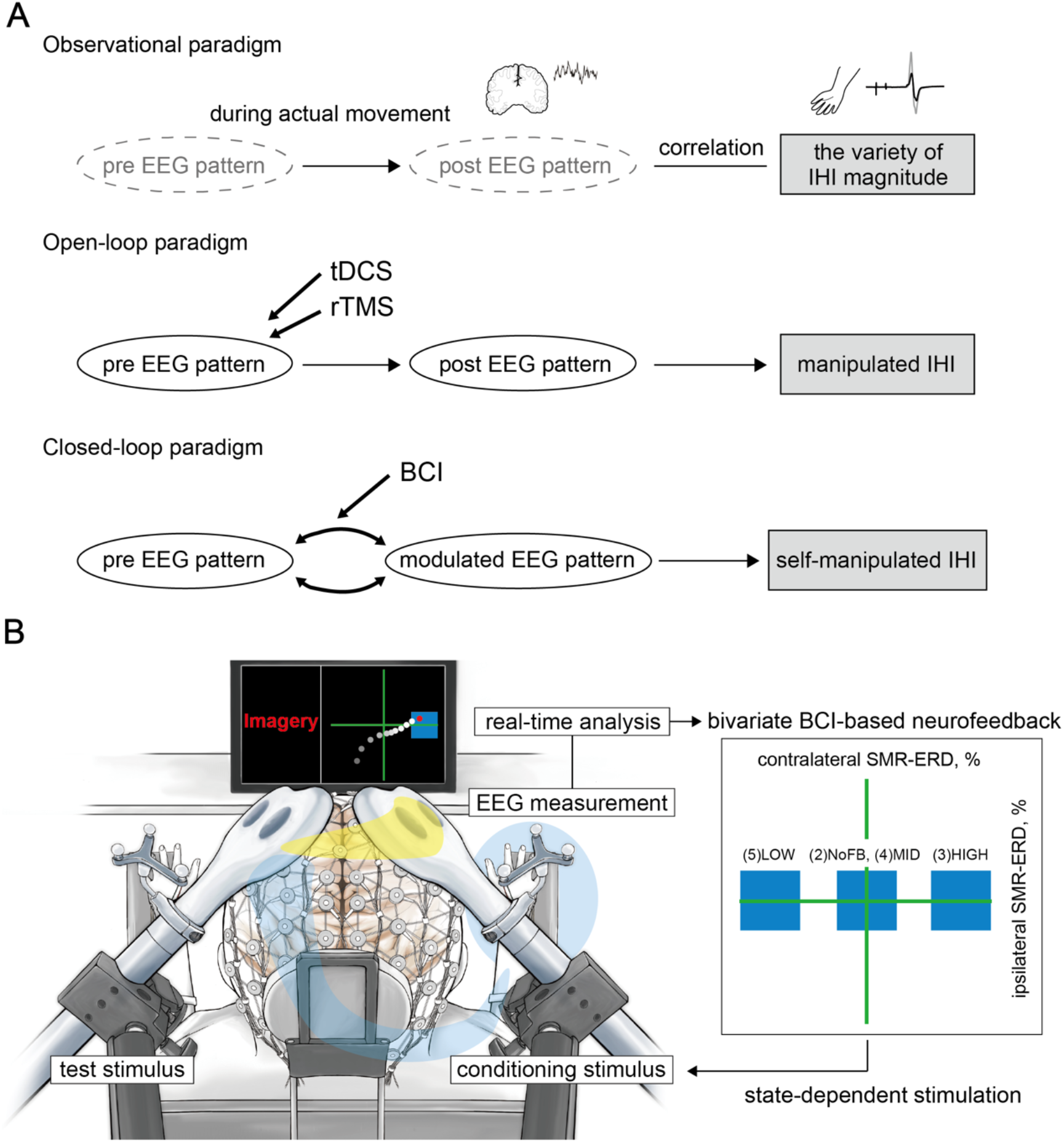
Conceptual illustration of the current study and experimental overview. **(A)** When a certain stimulus was input into the system, the brain was considered to vary with the state, resulting in IHI changes. The upper panel highlights the experimental limitations in the observational paradigm due to a variety of IHI magnitudes is observed during actual movement. In this case, it is unclear whether the changes in EEG patterns in both hemispheres would affect IHI. The middle panel indicates that it is unclear whether it is possible to consciously self-manipulate inhibitory interhemispheric sensorimotor activity in the open-loop neuromodulation paradigm using tDCS or rTMS. The lower panel shows that specific EEG patterns are associated with IHI magnitude, and closed-loop BCI-based neurofeedback modulates the EEG activities. Therefore, if bilateral EEG patterns that underlie IHI are identified, we should be able to volitionally up- and down-regulate the IHI magnitude via BCI-based neurofeedback, suggesting the possibility of plastic interhemispheric balancing. **(B)** The current bi-EEG-triggered dual-TMS experimental system involved spatially bivariate BCI-based neurofeedback that allows volitional modulation of EEG patterns only in a targeted hemisphere to enable us to verify our hypothesis. Different states of the targeted bidirectional up- and down-conditioned ipsilateral hemisphere to the imagined hand were tested in the following states: (1) resting state, (2) during motor imagery without visual feedback, (3) high, (4) middle, and (5) low excitability states. A blue target box, based on the predetermined SMR-ERDs, was displayed corresponding to each session. A cue signal was generated to trigger the conditioning stimulus when the signal reached the target box. The yellow line on the head represents the signal flow from the conditioning hemisphere that modifies the contralateral side through the corpus callosum, and the blue line represents the test signal towards the right hand.

As a recent neural manipulative tool, numerous studies have used a closed-loop brain-computer interface (BCI)-based neurofeedback whereby participants learn to volitionally desynchronize and synchronize oscillatory sensorimotor rhythms (SMR-ERD/ERS) in the contralateral sensorimotor cortex (SM1) through visual and/or somatosensory feedback (Ang and Guan, 2017; Chaudhary et al., 2016; Ramos-Murguialday et al., 2013; Soekadar et al., 2015b). BCI-based neurofeedback is supported by the fact that the intensity of SMR-ERD in EEG represents states of high versus low excitability of not only the sensorimotor cortex (Neuper et al., 2006; Neuper and Pfurtscheller, 2001; Pfurtscheller et al., 2006), but also the corticospinal descending pathway, as measured by the motor-evoked potential (MEP) amplitude (Takemi et al., 2015, 2013). Because both hemispheres are structurally and functionally connected, it is likely that the balance of bilateral SMR-ERDs and transcallosal excitability states are linked, as indicated by common variation of conditioning MEP amplitude and IHI (Ferbert et al., 1992; Ghosh et al., 2013; Ni et al., 2009).

Here, we sought to investigate the association of IHI and bilateral SMR-ERDs to understand the inhibitory sensorimotor functions of interhemispheric interaction that may critically depend on the oscillatory brain activity in both hemispheres. Furthermore, if the oscillatory brain activity reflected IHI, we tested whether it can be self-manipulated using a dual-coil paired-pulse TMS protocol (Daskalakis et al., 2002; Ferbert et al., 1992). To assess the association of IHI with ongoing oscillatory brain activity, we used a recently developed spatially bivariate BCI-based neurofeedback technique that allows volitional modulation of SMR amplitude in both hemispheres (Hayashi et al., 2021, 2020) and triggers the TMS pulses in real time at pre-specified bilateral SMR amplitudes. Our system, therefore, overcomes previously unresolved experimental limitations; unlike the ordinary observational or open-loop experimental paradigm, our paradigm enables participants to control the bivariate sensorimotor excitability, evaluates the inhibitory sensorimotor functions that underlie IHI, and determines whether it is possible to consciously self-manipulate the IHI magnitude.

Using the novel closed-loop bi-hemispheric brain state-dependent EEG-triggered dual-coil TMS system (bi-EEG-triggered dual-TMS), we evaluated the effects of different states of the targeted bidirectional up- or down-conditioned one-sided hemisphere on effective inhibitory interhemispheric network expressed by IHI: (1) resting state, (2) during motor imagery without visual feedback, (3) high, (4) middle, and (5) low excitability states of the ipsilateral SM1 (conditioning side) to the unilateral imagined hand movement during BCI-based neurofeedback (Figure 1B). We hypothesized that if the ipsilateral dominant SMR-ERD is a potent up- or down-regulator of IHI from the ipsilateral side, the strongest IHI will occur with high excitability state of the ipsilateral SM1. This multimodal research provides strong evidence for the dynamic interplay between distinct regions underlying IHI through BCI-based neurofeedback, and may therefore form the basis for self-manipulating IHI balancing to modulate the brain functions.

## Results

The data compliance is described in the Supplementary files (“Data compliance” and “IHI curves”).

### IHI manipulation via spatially bivariate BCI-based neurofeedback

We compared the IHI magnitude between five sessions as follows: (1) resting-state (REST); (2) right finger motor imagery (MI) without visual feedback (NoFB), (3) high excitability states of the ipsilateral SM1 during BCI-based neurofeedback (HIGH); (4) middle excitability states of the ipsilateral SM1 during BCI-based neurofeedback (MID); and (5) low excitability states of the ipsilateral SM1 during BCI-based neurofeedback (LOW). Typical examples of MEP amplitude elicited by single test stimulus (TS-only) and paired-pulse stimulation (conditioning stimulus [CS]+TS) of a representative participant are shown in Figure 2A. The manipulation range of IHI (i.e., the difference between HIGH and LOW sessions) was 32.6 ± 30.7% (Cohen’s *d* = 1.50). A one-way repeated-measures ANOVA (rmANOVA) of the sessions (five levels: REST, NoFB, HIGH, MID, and LOW) revealed significant differences (F_(4,88)_ = 6.85, *p* < 0.001, *η*^2^ = 0.22; REST: 69.0 ± 22.6%, NoFB: 85.9 ± 18.3%, HIGH: 62.0 ± 23.5%, MID: 85.5 ± 26.2%, LOW: 96.9 ± 27.7%). Across the three BCI-based neurofeedback sessions (i.e., HIGH, MID, and LOW sessions) for the comparison of IHI magnitude, a post-hoc two-tailed paired t-test showed significant difference between HIGH and MID sessions (difference = 23.5, Cohen’s *d* = 0.94, *p* = 0.025), and between HIGH and LOW sessions (difference = 34.9, Cohen’s *d* = 1.36, *p* = 0.001), but not between MID and LOW sessions (difference = 11.4, Cohen’s *d* = 0.42, *p* = 0.424; Figure 2B). Additional results are presented in Figure 2B. Importantly, rmANOVA for TS-only revealed no significant differences in the MEP amplitude between all sessions (F_(4,88)_ = 2.44, *p* = 0.104, *η*^2^ = 0.09; REST: 0.98 ± 0.45 mV, NoFB: 1.50 ± 1.02 mV, HIGH: 1.94 ± 1.37 mV, MID: 1.67 ± 1.15 mV, LOW: 1.93 ± 1.41 mV), suggesting that participants could learn to volitionally increase or decrease (bidirectional) the ipsilateral sensorimotor excitability while maintaining constant contralateral sensorimotor excitability (Figure 2C).

**Figure 2.**
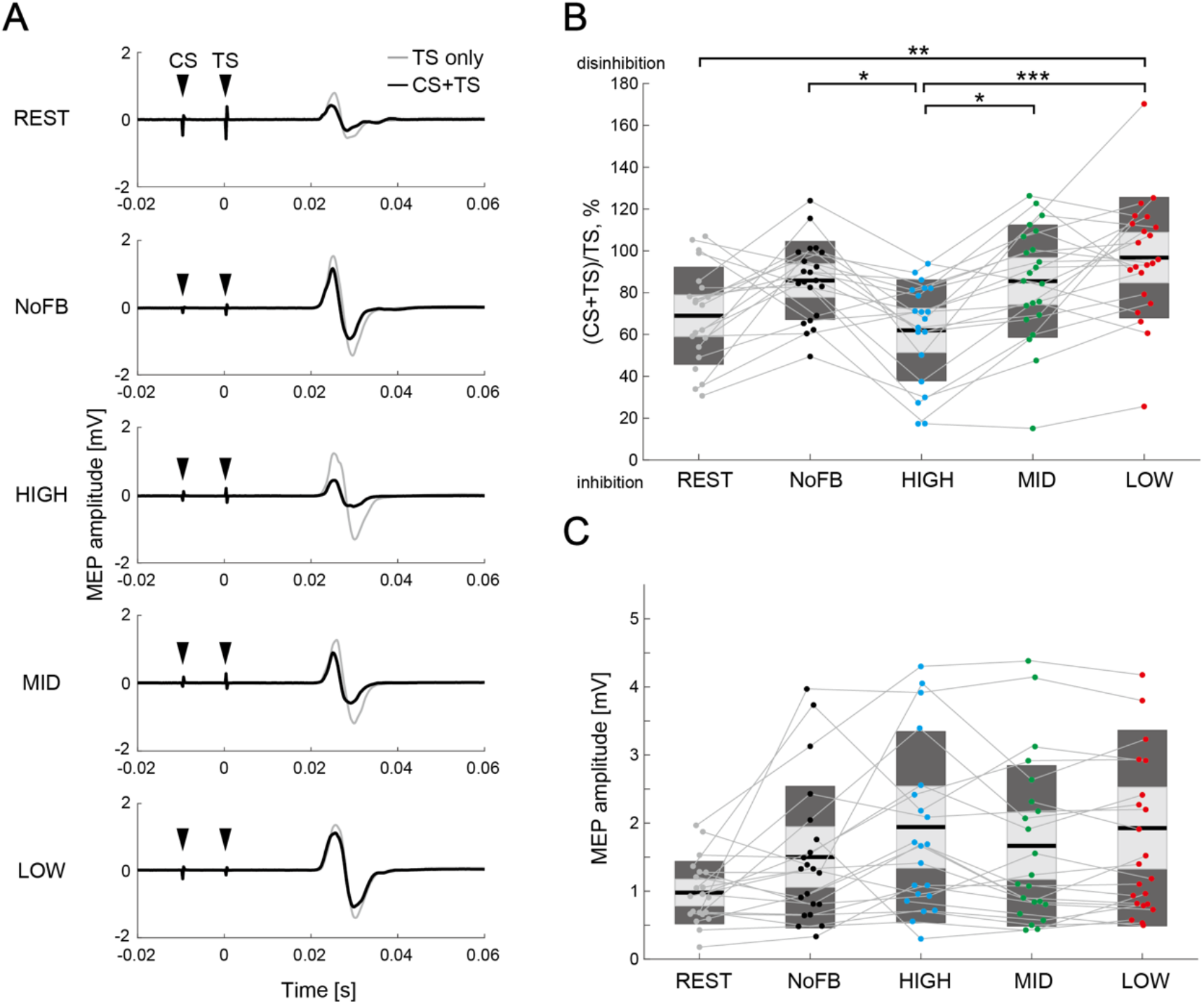
Comparison of IHI magnitude. **(A)** Typical examples of mean MEP amplitudes elicited by single TS (TS-only; light grey color) and paired-pulse stimulation (CS+TS; black color) in a representative participant. The black arrows represent the stimulus timings of CS and TS. **(B)** The IHI magnitude of the individual participants are represented by colored plots and thin grey lines. The figure also provides individual data in addition to the box plot. The light grey box represents 1.96 SEM (95% confidence interval) and dark grey box represents 1 SD. The black line indicates the group mean of the studied sample and colored plots represent a single session. Lower values represent greater inhibitory effect from the ipsilateral hemisphere and therefore greater acceleration of the IHI magnitude. Dendrograms above the bars represent the results of the post-hoc analyses. * *p* < 0.05, ** *p* < 0.01, and *** *p* < 0.001; all comparisons were Bonferroni corrected. **(C)** The figure shows MEP amplitude elicited by a single TS (TS-only). No significant difference in MEP amplitude was observed between sessions (all *p* > 0.05).

In the control muscle (abductor digiti minimi, ADM), such observed modulation was not driven (“IHI manipulation for control muscle” in Supplementary files). A linear mixed-effect model that considered the contralateral SMR-ERD and ipsilateral SMR-ERD as fixed effect, and participants as random effect revealed a significant effect of the ipsilateral SMR-ERD (*p* < 0.001) on IHI magnitude, but no main effect of the contralateral SMR-ERD or contralateral SMR-ERD ×ipsilateral SMR-ERD interaction (*p* = 0.828 and *p* = 0.058, respectively). We further compared the IHI magnitude across sessions by normalizing IHI magnitude to baseline (i.e., NoFB session), and calculating the difference between REST, HIGH, MID, and LOW sessions. The results were compatible with those presented in Figure 2 (“Comparison of normalized IHI magnitude” in Supplementary files). The results of the non-triggered TMS trials are also shown in the Supplementary files (“Comparison of IHI magnitude in non-triggered TMS trials”) to present the influence of spontaneous SMR fluctuations on IHI.

### Modulation effects of bilateral SM1 at EEG level

We tested whether participants could learn volitional modulation in both hemispheres at the EEG level using a spatially bivariate BCI-based neurofeedback. For the contralateral SMR-ERD, a one-way rmANOVA for the sessions (five levels: REST, NoFB, HIGH, MID, and LOW) revealed a significant difference (F_(4,96)_ = 4.81, *p* = 0.014, *η*^2^ = 0.16; REST: -10.6 ± 11.6%, NoFB: 13.5 ± 29.5%, HIGH: 19.3 ± 26.5%, MID: 16.1 ± 24.1%, LOW: 13.3 ± 28.4%). A post-hoc two-tailed paired t-test demonstrated no significant differences in the contralateral SMR-ERD between the neurofeedback sessions (HIGH-MID: difference = –3.2, Cohen’s *d* = 0.13, *p* = 1.00; HIGH-LOW: difference = –6.0, Cohen’s *d* = 0.22, *p* = 1.00; MID-LOW: difference = –2.8, Cohen’s *d* = 0.11, *p* = 1.00), whereas significant differences were found between REST and other sessions (all *p* < 0.05; Figure 3A). In contrast, after showing significant differences in the ipsilateral SMR-ERD between sessions (F_(4,96)_ = 156.0, *p* < 0.001, *η*^2^ = 0.86; REST: –18.8 ± 12.2%, NoFB: –10.5 ± 34.0%, HIGH: 64.5 ± 14.0%, MID: 10.1 ± 20.3%, LOW: –156.3 ± 49.0%), a post-hoc two-tailed paired t-test showed that ipsilateral SMR-ERD increased during the HIGH session (HIGH-MID: difference = –54.4, Cohen’s *d* = 3.12, *p* < 0.001; HIGH-LOW: difference = –220.8, Cohen’s *d* = 6.13, *p* < 0.001) and decreased during the LOW session (MID-LOW: difference = –166.4, Cohen’s *d* = 4.44, *p* < 0.001), revealing a significant modulation effect compared to baseline sensorimotor endogenous activity (Figure 3B). To investigate whether self-modulated changes in bilateral SMR-ERDs were specific to the hemisphere targeted by spatially bivariate BCI-based neurofeedback, we analyzed the laterality index (LI) for each session. A one-way rmANOVA for the sessions (five levels: REST, NoFB, HIGH, MID, and LOW) revealed significant differences (F_(4,96)_ = 53.9, *p* < 0.001, *η*^2^ = 0.68; REST: –0.14 ± 0.34, NoFB: –0.28 ± 0.31, HIGH: 0.54 ± 0.27, MID: –0.09 ± 0.42, LOW: –0.89 ± 0.14). The post-hoc two-tailed paired t-tests demonstrated that LI increased during the HIGH session (HIGH-MID: difference = –0.63, Cohen’s *d* = 1.78, *p* < 0.001; HIGH-LOW: difference = –1.43, Cohen’s *d* = 6.65, *p* < 0.001) and decreased during the LOW session (MID-LOW: difference = –0.80, Cohen’s *d* = 2.56, *p* < 0.001), indicating that the balance of interhemispheric sensorimotor excitability was modulated (Figure 3C). The spatial patterns of SMR-ERD in each session are depicted in Figure 3D. We found that the SMR-ERDs were predominantly localized in bilateral parieto-temporal regions (around the C3 and C4 channels and their periphery), and strong ipsilateral SMR-ERD was observed in the HIGH session with constant contralateral SMR-ERD.

**Figure 3.**
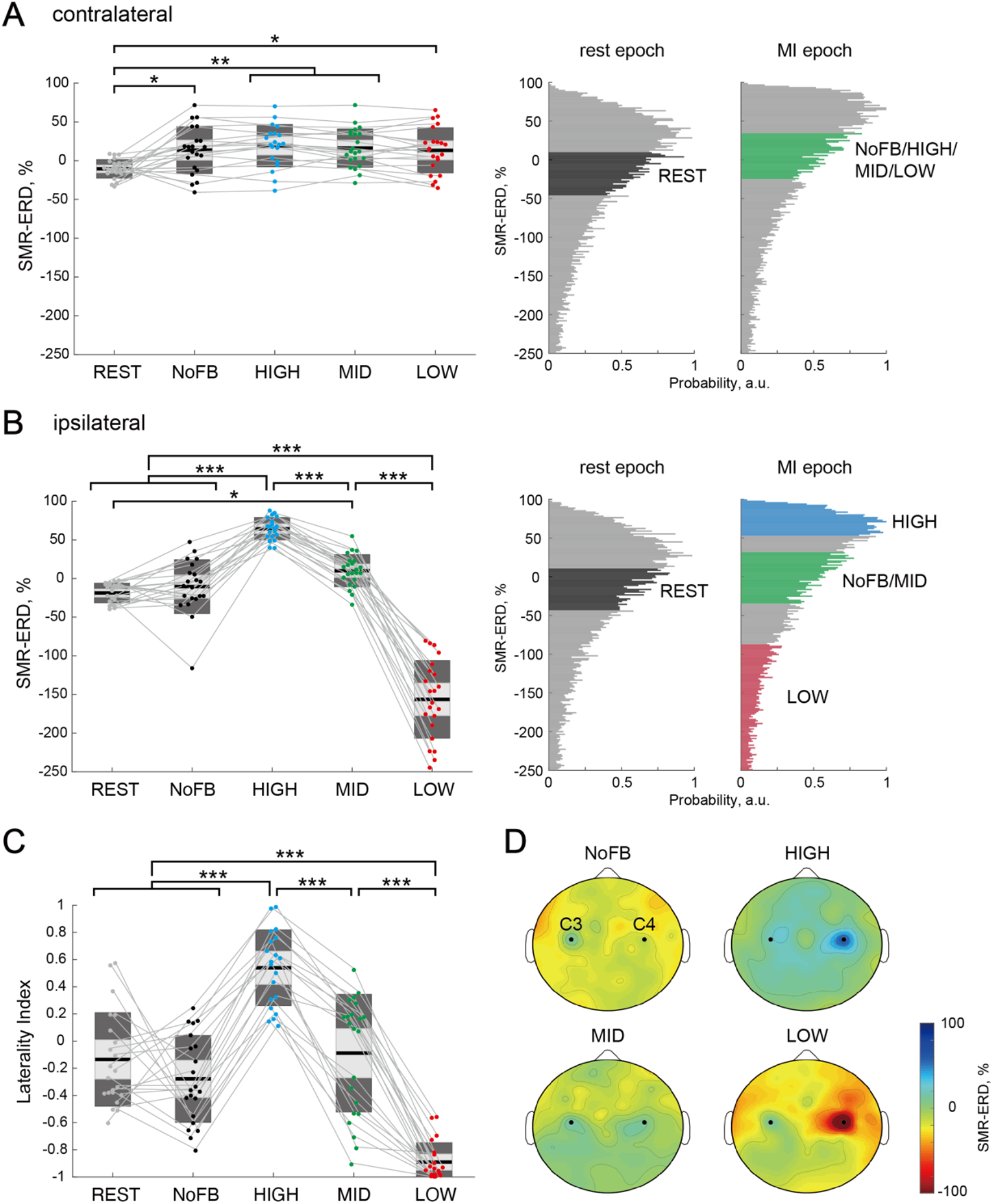
Target-hemisphere-specific modulation at the EEG level induced by spatially bivariate BCI-based neurofeedback. **(A), (B)** Modulation effect of the contralateral and ipsilateral SMR-ERDs, respectively. Individual participants are represented by colored plots and thin grey lines. The light grey box represents 1.96 SEM (95% confidence interval) and dark grey box represents 1 SD. The black line indicates the group mean of the studied sample and colored plots indicate a single session. Higher values represent greater sensorimotor excitability in the targeted hemisphere. The two right-sided panels represent the SMR-ERD distributions during rest and MI epoch in the calibration session. Based on the SMR-ERD distributions in the contralateral and ipsilateral hemispheres, the target ranges of SMR-ERD during bi-EEG-triggered dual-TMS system (each color) were set for each participant. **(C)** Laterality index (LI) for each session. LI yields a value of 1 or −1 when the activity is purely ipsilateral or contralateral, respectively. Dendrograms above the bars represent the results of the post-hoc analyses. * *p* < 0.05, ** *p* < 0.01, and *** *p* < 0.001; all comparisons were Bonferroni corrected. **(D)** Spatial patterns of SMR-ERD during the MI epoch in each session (group mean). Large positive values (blue color) represent larger SMR-ERD (i.e., higher excitability of the SM1). The black dots represent the C3 and C4 channels.

### Associations between IHI magnitude and bilateral EEG patterns

To examine the association between IHI magnitude and bilateral EEG patterns in each session, we performed a correlation analysis. In the within-subject correlation analysis between IHI magnitude and bilateral SMR-ERDs in each participant, 7 of the 22 participants showed a significant correlation between IHI magnitude and ipsilateral SMR-ERD (r = –0.587, *p* = 0.003 [representative]; r = –0.307 ± 0.252 [mean ± SD]), but not between IHI magnitude and contralateral SMR-ERD in three neurofeedback sessions (r = 0.020, *p* = 0.926 [representative]; r = –0.059 ± 0.298 [mean ± SD]; Figure 4A, B). In the across-subject correlation analysis between IHI magnitude and ipsilateral SMR-ERDs, we found a significant correlation by merging three neurofeedback sessions (r = – 0.330, *p* = 0.008), but not in either session (HIGH: r = –0.217, *p* = 0.359, MID: r = 0.037, *p* = 0.869, LOW: r = 0.022, *p* = 0.921; Figure 4C) due to inter-subject variability. The across-subject correlation analysis between IHI magnitude and contralateral SMR-ERDs showed no significant correlation by merging the three neurofeedback sessions (r = – 0.152, *p* = 0.231) or each session alone (HIGH: r = –0.079, *p* = 0.739, MID: r = –0.092, *p* = 0.685, LOW: r = –0.200, *p* = 0.377).

**Figure 4.**
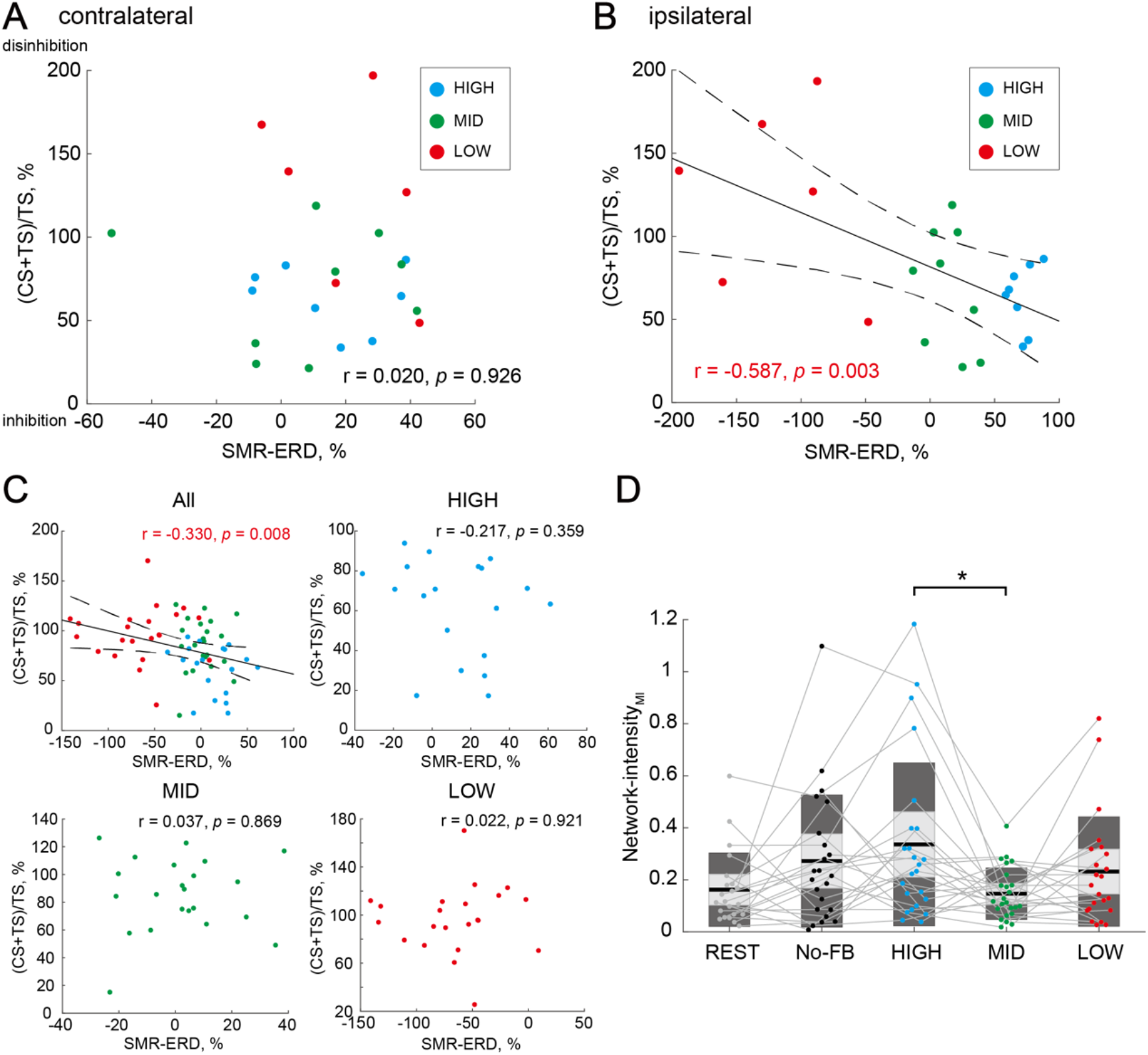
Associations of IHI magnitude and bilateral EEG patterns. **(A), (B)** Within-subject correlations between IHI magnitude and contralateral or ipsilateral SMR-ERDs in a representative participant. The plot colors indicate each session, with each plot representing a single triggered trial. Solid and dotted lines represent the estimated linear regression and 95% confidence interval, respectively. Only the ipsilateral SMR-ERD and IHI magnitude were significantly correlated; high sensorimotor excitability state in the ipsilateral hemisphere induced stronger inhibition from the ipsilateral hemisphere. **(C)** Across-subject correlations between IHI magnitude and ipsilateral SMR-ERDs in all three neurofeedback sessions (upper left), HIGH (upper right), MID (lower left), and LOW (lower right) sessions. Dots represent a single participant. No significant correlation was observed within either session. **(D)** Comparison of Network-intensity during MI across sessions. Dendrograms above the bars represent the results of the post-hoc analyses. * *p* < 0.05; all comparisons were Bonferroni corrected. There was a significant difference in interhemispheric functional connectivity between HIGH and MID sessions, but not between HIGH and LOW sessions, suggesting that interhemispheric functional connectivity did not reflect the excitatory or inhibitory activity.

To analyse the associations between IHI magnitude and bilateral neural network from another perspective, interhemispheric functional connectivity during MI was examined as a key EEG signature. For the Network-intensity_MI_, a one-way rmANOVA for the sessions (five levels: REST, NoFB, HIGH, MID, and LOW) revealed significant differences (F_(4,96)_ = 3.04, *p* = 0.020, *η*^2^ = 0.10; REST: 0.162 ± 0.138; NoFB: 0.272 ± 0.248; HIGH: 0.336 ± 0.306; MID: 0.147 ± 0.097; LOW: 0.232 ± 0.205). A post-hoc two-tailed paired t-test demonstrated significant differences between HIGH and MID sessions (difference = –0.189, Cohen’s *d* = 0.83, *p* = 0.033; Figure 4D). Surprisingly, however, we found no significant difference between HIGH and LOW sessions (difference = –0.104, Cohen’s *d* = 0.40, *p* = 1.00; Figure 4D), suggesting that interhemispheric functional connectivity did not reflect the excitatory or inhibitory activity.

### Distinct signatures for strong versus weak manipulation of IHI

Finally, we investigated the neural characteristics associated with the manipulation capability of IHI (ΔIHI_H-L_) using correlation-based analysis (Figure 5A). In the relationships between ΔIHI_H-L_ and IHI in the REST session (IHI_rest_), we found a significant correlation between ΔIHI_H-L_ and IHI_rest_ (r = –0.447, *p* = 0.044), indicating that participants with greater IHI at rest were able to strongly manipulate the IHI (Figure 5B). For the relationships between resting-state effective inhibitory interhemispheric network assessed by IHI_rest_ and interhemispheric functional connectivity at the EEG level (Network-intensity_rest_), participants with greater IHI_rest_ showed larger Network-intensity_rest_ (r = 0.547, *p* = 0.013; Figure 5C). Connectivity results in the other frequency bands are shown in Supplementary files (“Connectivity results in the other frequency bands”). Furthermore, we verified whether IHI_rest_ may be associated with intrinsic EEG profiles in the NoFB session including bilateral SMR-ERDs and LI during MI. We found a significant correlation between IHI_rest_ and ipsilateral SMR-ERD during MI (r = –0.619, *p* = 0.004), but not between IHI_rest_ and contralateral SMR-ERD during MI (r = –0.283, *p* = 0.228). Similarly, IHI_rest_ correlated with LI during MI (r = –0.457, *p* = 0.044; Figure 5D).

**Figure 5.**
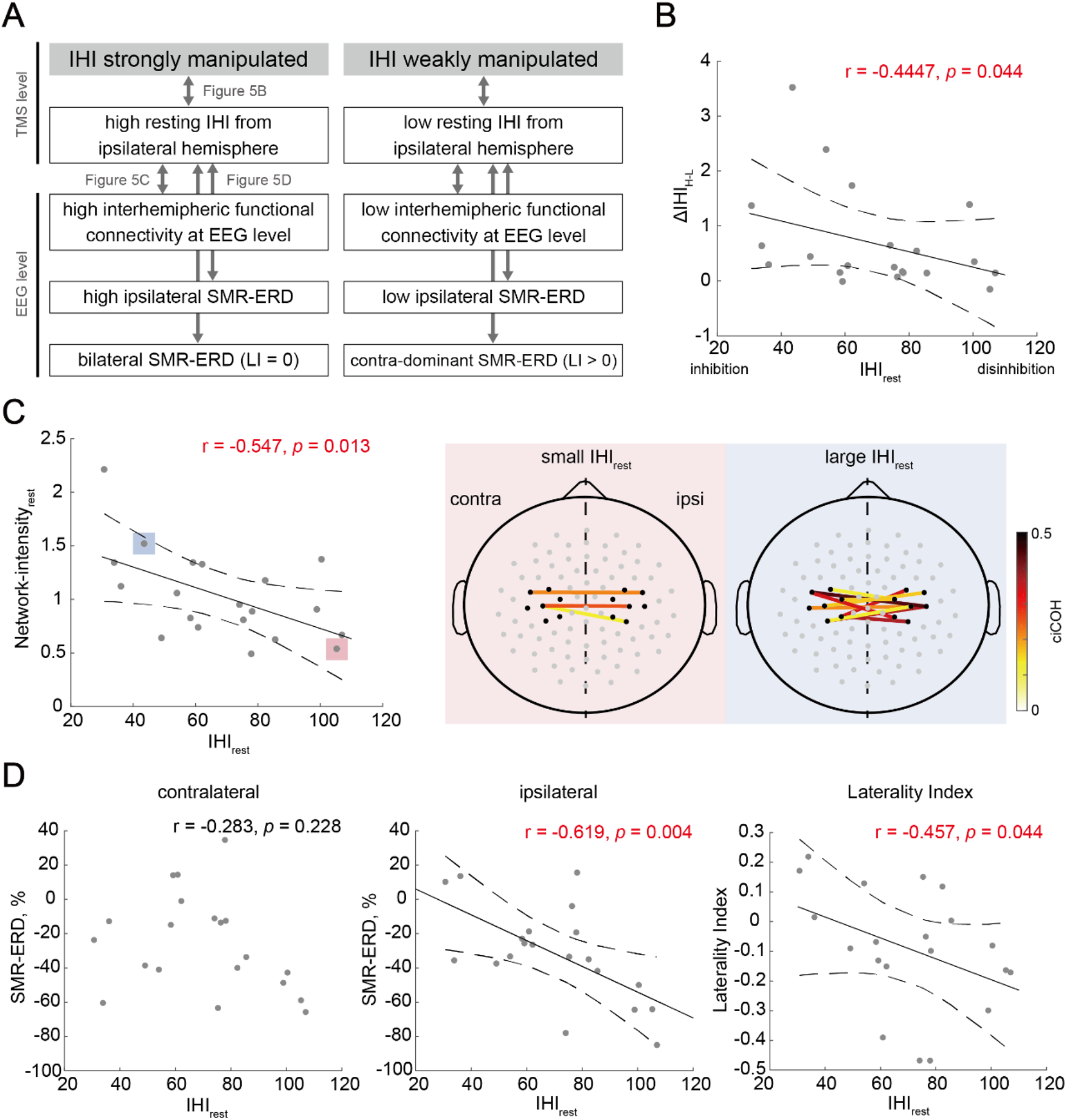
Neural characteristics depending on the manipulation capability of IHI. **(A)** Overview of the relationships between biomarkers from EEG and TMS data to probe the distinct signatures for strong versus weak manipulation of IHI. Arrows corresponds to a single panel. **(B)** Across-subject correlations between the manipulation capability of IHI and intrinsic IHI magnitude at rest. SMR-ERDs in a representative participant. Dots represent a single participant. Solid and dotted lines represent the estimated linear regression and 95% confidence interval, respectively. Participants with greater IHI at rest were able to strongly manipulate IHI. **(C)** Across-subject correlations between IHI at rest and EEG profiles showed significant correlations between IHI_rest_ and ipsilateral SMR-ERD and LI, but not between IHI_rest_ and contralateral SMR-ERD. **(D)** The left-sided panel shows across-subject correlations between IHI at rest and resting-state interhemispheric functional connectivity. The two right-sided panels indicate the significant interhemispheric connections (“Connectivity Analysis” in Methods) of the two representative participants with small and large IHI_rest_, respectively. The solid lines indicate a significant connection, and large positive values (dark red color) represent strong connections. The black dots around bilateral SM1 denote the seed channels, C3 or C4, and six neighboring channels. The gray dots represent other EEG channels.

## Discussion

In the present study, we aimed to determine whether it is possible to consciously self-manipulate the IHI magnitude and uncover related neural activity behind IHI, while controlling bilateral SMR-ERDs via closed-loop spatially bivariate BCI-based neurofeedback. This was the first study to show voluntary IHI-state manipulation with a large dynamic range related to variation in ipsilateral, rather than contralateral, sensorimotor excitability expressed by SMR-ERD and a resting-state interhemispheric network at the TMS (IHI_rest_) and EEG (interhemispheric Network-intensity_rest_) levels, which was accompanied by a balance of SMR-ERDs over bilateral SM1s.

Previous neurofeedback studies have demonstrated that it is possible to gain voluntary control over central nervous system activity without externally administered interventions, if appropriate neurofeedback is embedded in a reinforcement learning task, such as food rewards for animals (Engelhard et al., 2013; Fetz, 2013) and visually rewarding stimuli for humans (Thompson et al., 2009). However, conventional BCI-based neurofeedback of the SMR signal from one hemisphere does not guarantee spatial specific activation of the sensorimotor activity in the targeted hemisphere (Buch et al., 2008; Caria et al., 2011; Soekadar et al., 2015a) because sensorimotor activities in the left and right hemispheres potentially influence one another, making pathway-specific IHI manipulation difficult. Using spatially bivariate BCI-based neurofeedback enables participants to volitionally increase or decrease (bidirectional) the ipsilateral sensorimotor excitability, while maintaining constant contralateral sensorimotor excitability. Ipsilateral SMR-ERD (but not contralateral), reflecting IHI magnitude (Figure 3B) and their significant correlation (Figure 4C), were compatible with previous studies (Haegens et al., 2011; Madsen et al., 2019; Sauseng et al., 2009; Takemi et al., 2013; Thies et al., 2018; Zarkowski et al., 2006). The *in vivo* cortical recordings in monkeys revealed that pericentral alpha power was inversely related with the normalized firing rate in the sensorimotor regions (Haegens et al., 2011). In humans, several studies using single-pulse TMS of the motor hand area combined with EEG tested how ongoing pericentral oscillatory activity impacts corticomotor excitability reflected by the MEP amplitude. These EEG-TMS studies found a negative linear relationship between pre-stimulus SMR-ERD and MEP amplitude through both offline (Sauseng et al., 2009; Takemi et al., 2013; Zarkowski et al., 2006) and online (Madsen et al., 2019; Thies et al., 2018) approaches. As for IHI through the transcallosal fiber, it is predominantly regulated through direct postsynaptic mechanisms in the apical dendritic shafts of pyramidal neurons and a specific cortical microcircuitry mediated by dendritic GABA_B_ receptors in inhibitory interneurons (Palmer et al., 2012). Consistent with this phenomenon, down-regulation of ipsilateral SMR-ERD, for example, may influence the excitation of the ipsilateral transcallosal pyramidal neuron followed by disinhibition of the contralateral inhibitory interneurons during unilateral upper limb MI. Therefore, pronounced changes in cortical mechanism despite the absence of sensory input and constant MEP amplitude elicited by single TS suggest that the increase or decrease of IHI magnitude was at least partly of cortical, rather than spinal, origin. These neural mechanisms can be explored further by using triple pulse procedures (Ni et al., 2011) in which disinhibition can be directly measured as the reduction of short intracortical inhibition.

The potential contribution of resting-state IHI magnitude and interhemispheric functional connectivity in the alpha and high beta bands to the current results of manipulation capability in IHI (Figure 5B, C, and Supplementary figure 5C) can be speculated by considering the previous studies. Interregional communication is accompanied by synchronized oscillations in different brain regions (Fries, 2005; Varela et al., 2001), and this synchronization can be evaluated by functional connectivity. Because both hemispheres are structurally connected by transcallosal projection and exhibit functional cross talks (Hofer and Frahm, 2006; Meyer et al., 1995), the manipulation capability of IHI suggests it is associated with a structural connectivity (assessed by paired-pulse TMS and intrinsic EEG synchronization between hemispheres) and local oscillatory power entrainment. As for frequency band, it is well known that the intensity of SMR-ERD in the beta band reflects the sensorimotor cortical excitability and cortico-muscular activation (Hussain et al., 2019; Schulz et al., 2014). This multimodal EEG-TMS approach proves that the bilateral alpha and beta activities observed in the current study served to regulate the inhibition and facilitation of inhibitory functional coupling of interhemispheric interaction over motor cortices.

In addition, real-time fMRI-based neurofeedback to a single region of interest (Sitaram et al., 2017; Weiskopf et al., 2004) or interregional functional connectivity (Liew et al., 2016; Pereira et al., 2019) can be used for volitional self-modulation of neuronal connectivity and could serve as a possible therapeutic tool for motor or cognitive training in diseases related to impaired interregional interaction. However, even fMRI-based functional connectivity neurofeedback with high spatial resolution, based on blood oxygenation level-dependent signal, cannot differentiate between excitatory and inhibitory activities between these regions. Indeed, our results indicated no significant difference in Network-intensity during MI between HIGH and LOW sessions (Figure 4D). This interesting result would be explained by the following hypothesis: In LOW session, for example, down-regulation of the ipsilateral excitability leads to temporal down-regulation of the contralateral excitability since both hemispheres influence each other, which results in synchronized interhemispheric oscillatory activities; however, the contralateral excitability remains at the baseline level due to BCI-based neurofeedback and the interhemispheric input from the ipsilateral hemisphere increases simultaneously, which affect the down-regulation of intracortical inhibition in the contralateral hemisphere (Takemi et al., 2013), while maintaining constant right finger MEP. In addition, disinhibition of the inhibitory interneurons in the contralateral hemisphere projecting transcallosal pyramidal neuron causes increased IHI to the ipsilateral hemisphere, leading to similar sum of bidirectional IHI between HIGH and LOW sessions. Therefore, functional connectivity does not change. Hence, our results provided evidence that interhemispheric functional connectivity using fMRI cannot distinguish between excitatory and inhibitory neural activities, and ensured that our approach is useful as a direct IHI manipulator.

Interhemispheric activity is also passively modulated by externally administered interventions, for example, tDCS or rTMS over the motor cortices (Gilio et al., 2003; Peña-Gómez et al., 2012; Williams et al., 2010). Although such tools have neuromodulation efficacy, their long-term sustained effects are often limited and they result in local and remote effects without spatial specificity (Di Pino et al., 2014; Notturno et al., 2014; Weiskopf et al., 2004). Conversely, under the conscious self-learning environment, volitional control over MEP amplitudes is retained for at least 6 months without further training (Ruddy et al., 2018), which supports the prediction of long-term efficacy and safety of pathway-specific IHI self-manipulation. Furthermore, the effect size of IHI manipulation in the current study (Cohen’s *d* = 1.50) was approximately 1–2-fold higher than that of representative tDCS (Cohen’s *d* = 1.55) (Williams et al., 2010) and rTMS studies (Gilio et al., 2003) (Cohen’s *d* = 0.80; note that it is a read from the graph). Based on a previous study of neurofeedback training combined with an external administered interventions (Ang et al., 2012), their combination may be useful for facilitating neural plasticity.

Our evidence and techniques are expected to be applied in various fields, for example, in the context of neurorehabilitation. In stroke patients, the interhemispheric imbalance model predicts the presence of asymmetry in the interhemispheric sensorimotor network, with excessive inhibition from the non-affected hemisphere limiting maximal recovery (Murase et al., 2004). Therefore, guiding effective inhibitory interhemispheric network represented by IHI to the appropriate pattern through both targeted up-conditioning in the affected hemisphere and down-conditioning in the non-affected hemisphere may contribute to reduced abnormal IHI and enhanced achievable functional recovery (Chieffo et al., 2013; Di Pino et al., 2014; Dong et al., 2006; Hummel and Cohen, 2006). Although the rehabilitation strategy of attempting to rebalance interhemispheric interactions in order to improve motor recovery after stroke is controversial (Bundy et al., 2017; Xu et al., 2019), the current technique can be tailored either to upregulate the damaged hemisphere, down-regulate the intact hemisphere, a combination of both, or vice versa depending on the patient’s specific needs. Current proof-of-concept data reveal for the first time that self-manipulation of IHI via BCI-based neurofeedback is equal or superior to the conventional tools. Future work probing the residual ability to manipulate IHI in stroke patients is warranted.

In conclusion, we presented an innovative approach to voluntarily and bidirectionally manipulate the state of IHI, by directly targeting SMR-ERDs in a spatially bivariate BCI-based neurofeedback paradigm. This approach provides the opportunity to understand the inhibitory sensorimotor functions and paves the way for new technologies that allow the user/patient to regulate aspects of their brain function to reach the desired states, e.g., for neurorehabilitation and enhanced motor performance.

## Materials and Methods

### Study design

The current study was performed in accordance with approved guidelines and regulations, such as the CONSORT Statement (Moher et al., 2001) and CRED-nf checklist (Ros et al., 2020). The experiment consisted of five sessions: (1) resting-state (REST); (2) right finger MI without visual feedback (NoFB), (3) high (HIGH); (4) middle (MID); and (5) low excitability states of the ipsilateral SM1 during BCI-based neurofeedback (LOW) (details in “Experimental sessions”). The difference of IHI magnitude in the last three sessions (i.e., HIGH, MID, and LOW sessions) was the primary dependent variable of interest. REST and NoFB sessions were treated to estimate individual baseline during rest and MI for the offline analysis.

To estimate the appropriate sample size for this study, a preliminary experiment was conducted before the main experiment. In the preliminary experiment, four healthy participants (not included in the main experiment) performed BCI-based neurofeedback training and underwent brain state-dependent dual-coil brain stimulation, similar to the main experiment. We calculated the IHI magnitude in each session. Then, an a priori power analysis (α = 0.05, 1-β = 0.8, two-sided tests, Bonferroni corrected) focusing on the IHI magnitude using the statistical package G*Power 3.1 (Faul et al., 2009) was conducted. Because the preliminary experiment showed a large effect size on the IHI differences between HIGH (65.0 ± 22.8, mean ± SD) and MID (90.7 ± 23.4) sessions (Cohen’s *d* = 1.12), and between MID (90.7 ± 23.4) and LOW (106.3 ± 13.1) sessions (Cohen’s *d* = 0.82), we calculated that 24 participants were needed (Cohen, 2013, 1992).

### Participants

Twenty-four volunteers (2 females and 22 males; mean age ± SD: 23.4 ± 2.0 years; age range: 21–27 years) participated in this study. All participants had normal or corrected-to-normal vision and reported no history of neurological or psychological disorders. All participants were right-handed (Laterality Quotient: 72.2 ± 30.9%) as assessed by the Edinburgh Inventory (Oldfield, 1971). Participants were excluded if the resting motor threshold (RMT) of the right and left first dorsal interosseous (FDI) muscles was ≤ 70% of the maximum stimulator output (MSO) during the TMS experiment. This criterion ensured that the TMS stimulator would be able to perform at the required intensities for the whole duration of the experiment (Stefanou et al., 2018; Zrenner et al., 2018). We did not exclude participants based on the EEG characteristics such as the magnitude of their endogenous SMR activity (Madsen et al., 2019; Safeldt et al., 2017), to verify our hypothesis in a Proof-of-Concept study. Twenty-two participants (2 females and 20 males; mean age ± SD: 23.3 ± 1.9 years; age range: 21–27 years; Laterality Quotient: 72.5 ± 32.2%) attended the ensuing neurofeedback training with brain stimulation. The results presented in the EEG part are from all 24 participants, while IHI results are from the 22 participants that completed the whole experiment. Four of the 120 sessions from four participants (two REST, one NoFB, and one HIGH session) were excluded due to corrupted data.

The experiments conformed to the Declaration of Helsinki and were performed in accordance with the current TMS safety guidelines of the International Federation of Clinical Neurophysiology (Rossi et al., 2009). The experimental procedure was approved by the Ethics Committee of the Faculty of Science and Technology, Keio University (no.: 31-89, 2020-38, and 2021-74). Written informed consent was obtained from participants prior to the experiments.

### EEG/EMG data acquisition

EEG signals were acquired using a 128-channel Hydrogel Geodesic Sensor Net 130 system (Electrical Geodesics Incorporated [EGI], Eugene, OR, USA) in a quiet room. EEG data were collected at a sampling rate of 1 kHz and transmitted via an Ethernet switch (Gigabit Web Smart Switch; Black Box, Pennsylvania, USA) to the EEG recording software (Net Station 5.2; EGI and MATLAB R2019a; The Mathworks, Inc, Massachusetts, USA). The ground and reference channels were placed at CPz and Cz, respectively. The impedance of all channels, excluding the outermost part, was maintained below 30 kΩ throughout the experiment to standardize the EEG recordings (Ferree et al., 2001). This impedance standard was consistent with other studies using the same EEG system (Carter Leno et al., 2018b, 2018a; Robertson et al., 2019).

Surface EMGs were recorded from the FDI and ADM of the left and right hands using two pairs of Ag/AgCl electrodes (*ϕ* = 10 mm) in a belly-tendon montage. Impedance for all channels was maintained below 20 kΩ throughout the experiment. EMG signals were digitized at 10 kHz using Neuropack MEB-2306 (Nihon Kohden, Tokyo, Japan). The EMG data from each trial were stored for offline analysis on a computer from 500 ms before to 500 ms after the TMS pulse. Simultaneously, 5–10 ms of data were transferred immediately after collection to a computer for real-time analysis. In case of muscular contraction due to finger movement, the experimenter reminded participants to relax their muscles and ensure absence of muscle activity during MI. To monitor the real-time surface EMG signals, EMG signals were band-pass filtered (5– 1000 Hz with 2nd order Butterworth) with a 50-Hz notch to avoid power-line noise contamination; the root mean square of the filtered EMG signal from the FDI for the previous 1000 ms of data was displayed on the second experimenter’s screen in the form of a bar.

Throughout the experiment, the participants were seated in a comfortable chair with stable forearm support and performed unilateral right index finger abduction MI. The wrist and elbow joint angles were fixed to a neutral posture at the armrest. The participants were instructed to maintain this posture and were visually monitored by the experimenter throughout the EEG and MEP measurements. During MI, the forelimbs were placed in a prone position, with natural elbow and shoulder joint angles to help prevent the muscle activity.

### TMS protocol

For the TMS experiments, TMS was delivered using two interconnected single-pulse magnetic stimulators (The Magstim BiStim^2^; Magstim, Whiteland, UK) producing two monophasic current waveforms in a 70-mm figure-of-eight coil. We identified the optimal left and right coil positions over the hand representation area at which a single-pulse TMS evoked a MEP response in the FDI muscle with the lowest stimulus intensity, referred to as the motor hotspot. The TS was delivered to the motor hotspot of the right M1, with the handle of the coil pointing backward and approximately 45° to the midsagittal line. The other coil for the CS was similarly placed over the motor hotspot of the left M1 but slightly reoriented at 45–60° relative to the midsagittal line because it was not possible to place two coils in some participants with small head size. This orientation is often chosen in IHI studies (Daskalakis et al., 2002) since it induces a posterior-anterior current flow approximately perpendicular to the anterior wall of the central sulcus, which evokes MEPs at the lowest stimulus intensities (Rossini et al., 2015). To immobilize the head and maintain fixed coil positions over the motor hotspots during the experiment, chin support and coil fixation arms were used. The position of the TMS coil was monitored using the Brainsight TMS navigation system (Rogue Research, Cardiff, UK), so that the optimal coil orientation and location remained constant throughout the experiment.

RMT was defined as the lowest stimulator output eliciting an MEP in the contralateral side of relaxed FDI of > 50 μV peak-to-peak in 5 out of 10 consecutive trials (Groppa et al., 2012; Rossini et al., 1994). The stimulus intensities of the left and right M1 to evoke MEP of 1 mV peak-to-peak amplitude from the relaxed right and left FDIs (SI_1mV_) were also determined for the following dual-coil paired-pulse TMS. The average SI_1mV_ of TS and CS were 124 ± 11% RMT and 132 ± 13% RMT, respectively.

### IHI evaluation

IHI from the ipsilateral (right) to the contralateral (left) M1 was probed using a dual-coil paired-pulse TMS paradigm. CS was applied to the right M1, followed a few milliseconds later by a TS delivered to the left M1 (Ferbert et al., 1992). Due to the time constraint, the ISI in the present study was uniformly set to 10 ms, in accordance with previous studies (Duque et al., 2005; Harris-Love et al., 2007; Murase et al., 2004; Tsutsumi et al., 2012); however, an ideal ISI would vary across individuals. Additionally, in a preliminary experiment with four participants, it was confirmed that IHI was clearly observed when ISI was set to 10 ms. The stimulus intensity remained constant throughout the experiment for each participant.

To validate IHI measurement under bi-EEG-triggered dual-TMS setup, IHI curves were obtained in 20 of 24 participants prior to the main experiment, where a CS of varying intensity (five different intensities, 100–140% of RMT, in steps of 10% RMT) preceded the TS. Ten conditioned MEPs were collected for each CS intensity, along with 10 unconditioned MEPs (i.e., TS was given alone) in randomized order. The peak-to-peak amplitudes of the conditioned MEPs were averaged for the different CS intensities and expressed as a percentage of the mean unconditioned MEP amplitude. IHI intensity curves (Figure 2) ensured that IHI was approximately half-maximum for each participant when 120-130% RMT of CS intensity was applied, similar to previous EEG-TMS experiment (Stefanou et al., 2018; Tsutsumi et al., 2012).

### Spatially bivariate BCI-based neurofeedback

The present study was conducted based on a spatially bivariate BCI-based neurofeedback that displays bi-hemispheric sensorimotor cortical activities, which we recently developed in our laboratory (Hayashi et al., 2021, 2020). This method allows participants to learn to regulate these two variates at the same time and induce changes in target-hemisphere-specific SMR-ERD. Visual feedback was provided on a computer screen in the form of cursor movements in a two-dimensional coordinate, in which x and y axis corresponded to the degree of the ipsilateral and contralateral SMR-ERD, respectively. The axis range was set from the 5th (i.e., SMR-ERS) to 95th (i.e., SMR-ERD) percentile of intrinsic SMR-ERD distribution in EEG calibration session, and the origin-position (x= 0, y = 0) represented median values of bilateral SMR-ERDs. The cursors were presented at the origin-position at the initiation of a trial, and values exceeding the boundary were rounded to the 5th or 95th percentile. A key point of the current methodology is that, for example, when participants were instructed to move the cursor toward the middle right (x > 0, y = 0) in the two-dimensional coordinate, the position showed a strong SMR-ERD in the ipsilateral hemisphere and moderate SMR-ERD in the contralateral hemisphere.

Therefore, spatially bivariate BCI-based neurofeedback enables us to investigate how sensorimotor excitability in the target hemisphere (i.e., ipsilateral side) contributes to IHI while maintaining constant contralateral sensorimotor excitability.

During MI, participants were asked to perform a right index finger kinesthetic MI from the first-person perspective with equal time constants of 0.5 Hz cycle. Kinesthetic MI was performed because a previous study demonstrated that the focus of EEG activity during kinesthetic MI was close to the sensorimotor hand area, whereas visual MI did not reveal a clear spatial pattern (Neuper et al., 2005). To improve MI task compliance (i.e., whether all participants successfully performed the MI in the same manner), we not only asked them to perform kinesthetic MI from a first-person perspective, but also asked them to perform a rehearsal before each session. In addition, we confirmed that SMR was observed in a frequency-specific, spatiotemporal-specific, and task-related manner through offline analysis after each session. These characteristics of SMR-ERD indicate that kinesthetic MI, not visual MI, was performed correctly (Neuper et al., 2005; Pfurtscheller and Neuper, 1997).

### Real-time brain state-dependent dual-coil brain stimulation

The bi-EEG-triggered dual-TMS setup uses an online output of the raw EEG signal and analyzes it in real time to trigger TMS pulses depending on the instantaneous bilateral spatially filtered SMR-ERD of the recorded EEG. The real-time SMR-ERD intensity in each hemisphere (relative to the average power of the 1-5 s of the resting epoch) was obtained every 100 ms and calculated using the last 1-s data as follows (Hayashi et al., 2020): (1) acquired raw EEG signals recorded over SM1 underwent a 1–70-Hz second-order Butterworth bandpass filter and a 50-Hz notch filter; (2) filtered EEG signals were spatially filtered with a large Laplacian (60 mm to set of surrounding channels), which subtracted the average value of the surrounding six channel montage from that of the channel of interest (i.e., C3 and C4, respectively). This method enabled us to extract the task-related EEG signature and improve the signal-to-noise ratio of SMR signals (McFarland et al., 1997; Tsuchimoto et al., 2021). In addition, the large Laplacian method is better matched to the topographical extent of the EEG control signal than the small Laplacian and ear reference methods (McFarland et al., 1997); (3) a fast Fourier transform was applied to the spatially large Laplacian filtered EEG signals; (4) the power spectrum was calculated by calculating the square of the Fourier spectrum; (5) the alpha band power was obtained by averaging the power spectrum across the predefined alpha target frequencies from the EEG calibration session (described below); (6) the alpha band power was time-smoothed by averaging across the last five windows (i.e., 500 ms) to extract the low-frequency component for high controllability. The slow fluctuation component is beneficial to neurofeedback training because it reduced the flickering and improved the signal-to-noise ratio of the SMR signal (He et al., 2020; Kober et al., 2018); and (7) SMR-ERD was obtained by calculating the relative power to the average power during the resting epoch. SMR-ERDs of the last ten segments were displayed and updated every 100 ms, allowing participants to compare the current and past conditions. Thereafter, a cue signal was generated to trigger the magnetic stimulator of CS stimulus when the signal reached the predetermined target that the SMR-ERD threshold was exceeded and transmitted Transistor-Transistor Logic pulse to the magnetic stimulator of TS stimulus 10 ms later by Neuropack MEB-2306 system.

### Experimental sessions

Before the main IHI experiment, maximal voluntary contraction (MVC) was measured (Figure 6A). Full-length isometric abduction of the right and left index and little fingers were performed once after several exercises; each execution lasted 5 s with a 30-s rest between contractions to allow for recovery from mental fatigue. Each MVC was obtained by calculating the root mean square of stable 3 s of filtered EMG data.

**Figure 6.**
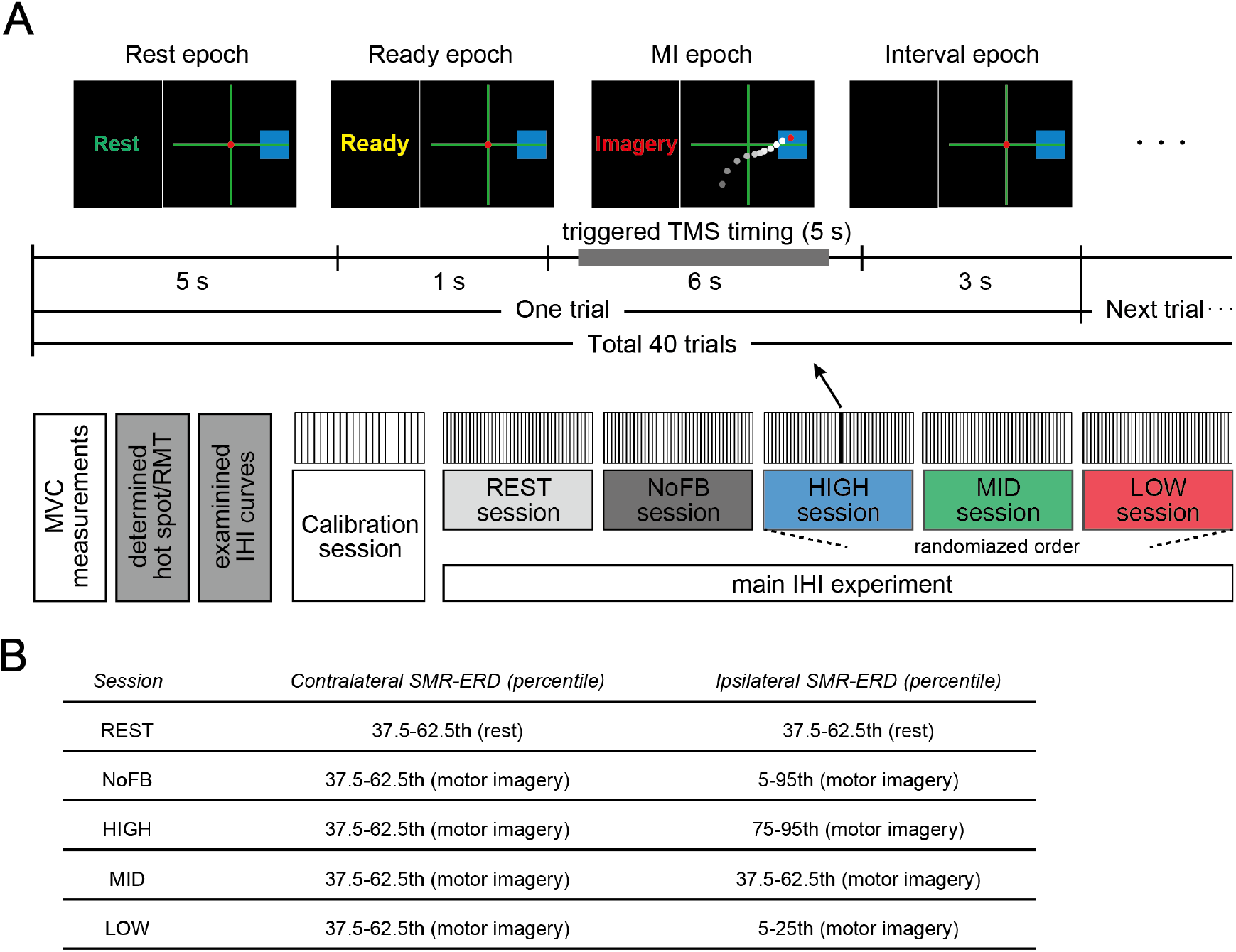
Experimental paradigm. **(A)** Task instructions and visual SMR-ERD feedback in the contralateral and ipsilateral SM1 were provided in the form of computer cursors in a two-dimensional coordinate on a computer screen (upper panel). EEG-triggered TMS timing ranged from 0.5 to 5.5 s during the MI epoch, and non-triggered TMS (referred to as the failed trial) was delivered in shuffled timing ranging from 5.5 to 6 s during the MI epoch to determine the influence of spontaneous SMR fluctuations on IHI. Lower panel indicates the experimental overview. The last three HIGH, MID, and LOW sessions were arranged in random order, and in these sessions, participants received visual feedback based on bilateral SMR-ERDs. **(B)** The predefined target ranges of SMR-ERD were expressed by a blue rectangle on the computer screen in each session. We aimed for participants to volitionally increase or decrease (bidirectional) the ipsilateral sensorimotor excitability while maintaining constant contralateral sensorimotor excitability. REST and NoFB sessions were served in order to estimate the individual baseline during rest and MI for the offline analysis.

Next, to determine the parameters of the bi-EEG-triggered dual-TMS setup, an EEG calibration session consisting of 20 trials, providing real-time SMR-ERD only on the contralateral side, was performed for each participant prior to the IHI experiment. Each trial was initiated by a 5-s resting epoch, followed by a 1-s ready epoch, and completed by a 6-s MI epoch. During this 12-s trial period, participants were asked not to move, blink, or swallow to prevent EEG artefacts derived from non-neural activity. After each 12-s trial, the screen went black for 3 s (Figure 6A). Participants were allowed to move freely to avoid mental fatigue during this interval period, before the next trial started. Thereafter, the target frequencies in the contralateral and ipsilateral SM1 were determined for each participant in order to feedback the most reactive frequency. Since SMR-ERD in the alpha band is a reliable EEG biomarker of increased neuronal excitability in SM1, corticospinal tract, and thalamocortical systems (Neuper et al., 2006; Soekadar et al., 2015b; Takemi et al., 2018, 2015, 2013; Yuan et al., 2010), the target frequencies were selected from the alpha band (8–13 Hz) by calculating the mean intensity of SMR-ERD with a 3-Hz sliding bin and 2-Hz overlap. Second, the target ranges of SMR-ERD during bi-EEG-triggered dual-TMS setting were normalized for each participant based on SMR-ERD distribution in the contralateral and ipsilateral hemispheres.

After the calibration session, the main IHI experiment with dual-coil paired-pulse TMS was performed in five consecutive sessions (10-min each) with fixed CS and TS intensities. Each session consisted of 40 trials and the task sequence of each trial was similar to that of the EEG calibration session. Five experimental sessions comprised different conditions as follows: (1) resting-state where participants were instructed to relax and look at the origin of the 2-D coordinates on the computer screen in front of them (REST); (2) right finger MI without visual feedback (NoFB); participants tried to achieve (3) high (HIGH); (4) middle (MID), and (5) low excitability states of the ipsilateral SM1 during BCI-base neurofeedback (LOW). The last three HIGH, MID, and LOW sessions were arranged in a random order, and in these sessions, participants received visual feedback based on the SMR-ERDs from both contralateral and ipsilateral hemispheres.

In each session, five trigger conditions were tested, with approximately equal numbers of paired pulses and unconditioned test pulses. EEG-triggered TMS timing ranged from 0.5 to 5.5 s during the MI epoch, and non-triggered TMS (referred to as the failed trial) was delivered in shuffled timing ranging from 5.5 to 6 s during the MI epoch to see the influence of spontaneous SMR fluctuations on IHI. Trigger conditions were determined based on the intrinsic sensorimotor cortical activity of each participant and was calculated in the EEG calibration session. The predetermined target ranges of SMR-ERD was expressed by a blue rectangle on the computer screen in each session are as follows (Figure 6B): (1) 37.5–62.5th percentile of SMR-ERD distribution during rest in both hemispheres (REST session); (2) 37.5–62.5th percentile of SMR-ERD distribution during MI in the contralateral hemisphere and 5–95th percentile of SMR-ERD distribution during MI in the ipsilateral hemisphere (NoFB session); (3) 37.5–62.5th percentile of SMR-ERD distribution during MI in the contralateral hemisphere and 75– 95th percentile of SMR-ERD distribution during MI in the ipsilateral hemisphere (HIGH session); (4) 37.5–62.5th percentile of SMR-ERD distribution during MI in the contralateral hemisphere and 37.5–62.5th percentile of SMR-ERD distribution during MI in the ipsilateral hemisphere (MID session); and (5) 37.5–62.5th percentile of SMR-ERD distribution during MI in the contralateral hemisphere and 5–25th percentile of SMR-ERD distribution during MI in the ipsilateral hemisphere (LOW session). The last three sessions (i.e., HIGH, MID, and LOW sessions) were our primary dependent variables of interest to verify how SMR-ERD in the ipsilateral hemisphere contributes to IHI while maintaining constant contralateral SMR-ERD. The REST and NoFB sessions served as controls to determine the intrinsic IHI magnitude without a neurofeedback paradigm. To evaluate the difficulty of each neurofeedback session, the mean of the sum of the success triggered trials (± 1 SD) of all sessions (except NoFB session because it has a wide triggered range) was calculated. The waiting time for a triggered event from MI onset was measured for each session.

### MEP analysis

For the quality control of MEP analysis, trials were rejected if: (1) coil position was shifted from the optimal orientation and location (> 3 mm and/or > 3°) despite maintaining it during the experiment using the Brainsight TMS navigation system; (2) involuntary muscle contraction in the 250 ms period before the TMS pulse was observed (> 5% MVC) because of pre-innervation increase in MEP amplitude (Devanne et al., 1997; Hallett, 2007); (3) large trial-by-trial MEP variance (mean ± 3SD) were found in order to screen out extreme values (Ruddy et al., 2018). In total, 8.8% of all trials were excluded from further analysis. Each peak-to-peak MEP amplitude was automatically determined in the remaining trials within 20–45 ms after the TMS pulse. IHI was defined as the percentage of mean conditioned MEP amplitude over mean unconditioned MEP amplitude (IHI = conditioned MEP/unconditioned MEP × 100%); therefore, smaller IHI values represent stronger inhibition. EMG and EEG data were processed using the customized analysis scripts on MATLAB R2019a.

### Offline EEG analysis

To evaluate the sensorimotor excitability that may influence IHI, pre-processing and time-frequency analyses were performed, and the left and right SMR-ERDs and their laterality were calculated. SMR-ERD in EEG is a reliable surrogate monitoring marker of sensorimotor excitability level for several reasons: (1) SMR-ERD and task-induced increase in blood oxygenation level-dependent signals during MI are co-localized and co-varied at SM1 (Yuan et al., 2010); (2) SMR-ERD control is associated with the contribution of SM1 modulated by transcranial direct current stimulation (Soekadar et al., 2015b); and (3) data-driven EEG features discriminating the presence or absence of muscle contraction were predominantly localized in the parieto-temporal regions, indicating SMR-ERD (Hayashi et al., 2019; Iwama et al., 2020). The time segment of interest was from the initiation of the trial to before TMS-triggered time marker of the CS in order to avoid contamination by the TMS artifact (pre-stimulation period). The EEG signal underwent a 1–70-Hz, second-order Butterworth bandpass filter and a 50-Hz notch filter. The EEG signals of all channels were spatially filtered using a common average reference, which subtracted the average value of the entire electrode montage (the common average) from that of the channel of interest to remove global noise (McFarland et al., 1997; Tsuchimoto et al., 2021). EEG channels in each trial were rejected during further analysis if they contained an amplitude above 100 μV (Sanei and Chambers, 2013). To examine target-hemisphere-dependent modulation (i.e., difference between the contralateral and ipsilateral sensorimotor activation), LI of the bihemispheric SMR-ERDs was calculated (Seghier, 2008). LI yields a value of 1 or –1 when the activity is purely ipsilateral or contralateral, respectively.

### Connectivity analysis

To assess interhemispheric functional connectivity at the EEG level, distributed interregional neural communication was calculated. We focused on both the resting epoch (1-5 s) and MI epoch (7 to before stimulation onset) for analysis. To calculate functional connectivity and compensate for long-range synchronization preference, we used the corrected imaginary part of coherence (ciCOH) (Ewald et al., 2012; Hayashi et al., 2020; Vukelić and Gharabaghi, 2015). The details of the following processing for the connectivity analysis can be obtained from our previous work (Hayashi et al., 2020). The ciCOH was obtained by subdividing the resting epoch into 1-s segments with 90% overlap (31 segments in total) and multiplied with a Hanning window. Then, interhemispheric Network-intensity was computed as follows:

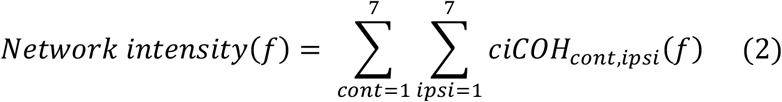

where *Network-intensity* is computed between left and right hemisphere, ∑ ∑ *ciCOH* is the sum of significant ciCOH values for interhemispheric interaction, *cont* denotes the 7 channels of interest (i.e., C3 and its neighboring 6 channels), *ipsi* denotes the 7 channels of interest in the opposite hemisphere (i.e., C4 and its neighboring 6 channels), and *f* indicates the frequency of interest (i.e., the predefined alpha frequencies). As a negative control, results of other frequency bands are shown in Supplementary files (“Connectivity results in the other frequency bands”).

### Correlation analysis

To investigate the association between sensorimotor brain activity at the EEG level and IHI magnitude from multimodal perspectives, correlation analysis was performed. We used Pearson’s correlation since EMG/EEG data showed normal distribution. Within-subject correlations between bilateral SMR-ERDs and IHI magnitude in each participant and across-subject correlations were performed. Furthermore, in order to examine the neural characteristics depending on the manipulation capability of IHI, we first examined the association between the manipulated effects on IHI calculated from the difference between HIGH and LOW sessions (ΔIHI_H-L_), and IHI in REST session (IHI_rest_). Next, the correlations between IHI_rest_ and intrinsic EEG profile in NoFB session (i.e., contralateral SMR-ERD, ipsilateral SMR-ERD, and LI) were investigated, respectively. Moreover, we verified whether large-scale resting-state functional connectivity was associated with an effective inhibitory interhemispheric network assessed by IHI. An across-subject Pearson’s correlation was applied to identify significant relationships between the IHI_rest_ and interhemispheric Network-intensity_rest_ for all participants

### Statistical analysis

Statistical analyses were performed using SPSS software (version 27; IBM Corp., Armonk, NY, USA) and MATLAB R2019a. The assumption of normality was verified using the Shapiro-Wilk test. All data were normally distributed (*p* > 0.05) and therefore analyzed with parametric tests. The assumption of sphericity was checked using the Mauchly’s test. If the test was significant, a Greenhouse-Geisser correction was applied. For the IHI curves, a one-way rmANOVA for intensities (six levels: 0% [TS only], 100%, 110%, 120%, 130%, and 140% RMT) and post-hoc two-tailed paired t-tests were performed in MEP amplitude. Modulation effects of bilateral sensorimotor excitabilities due to BCI-base neurofeedback training were evaluated from the 1-s period immediately before stimulation onset. A one-way rmANOVA for sessions (five levels: REST, NoFB, HIGH, MID, and LOW) was performed using the contralateral and ipsilateral SMR-ERDs and LI. Following the one-way rmANOVA, post-hoc two-tailed paired t-tests were performed using the Bonferroni correction for multiple comparisons. For the interhemispheric connectivity during MI, a one-way rmANOVA and post-hoc analysis were applied to Network-intensity_MI_ as the same procedure. In addition, to verify the difficulties for neurofeedback sessions, one-way rmANOVA for sessions (HIGH, MID, and LOW) and post-hoc analysis were applied to the number of the triggered trials and mean waiting time for a triggered event from MI onset. For the IHI results, a one-way rmANOVA for sessions (five levels: REST, NoFB, HIGH, MID, and LOW) was performed. Next, for all significant main effects, post-hoc two-tailed paired t-test was performed using the Bonferroni correction for multiple comparisons for all sessions. We further compared the IHI magnitude across sessions, by normalizing IHI magnitude to baseline (i.e., NoFB session) and calculating the difference between sessions (REST, HIGH, MID, and LOW). The significance level for all statistical tests was set to *p* = 0.05.

Although analysis of mean IHI magnitude across subjects revealed significant effect of the ipsilateral SMR-ERD (Figure 3B), we further performed mixed-effects analysis (Hussain et al., 2019; Madsen et al., 2019) incorporating the contralateral SMR-ERD and ipsilateral SMR-ERD as factors in the statistical model to explore the relationships between inhibitory interhemispheric activity and EEG characteristics. The linear mixed-effect model included the contralateral and ipsilateral SMR-ERD as fixed effects, treating the participant factor as a random effect to account for individual variability in IHI magnitude.

## Abbreviations

EEG: Electroencephalogram
TMS: transcranial magnetic stimulation
IHI: interhemispheric inhibition
BCI: brain-computer interface
SMR: sensorimotor rhythms
(ERD/ERS): event- related desynchronization/synchronization
SM1: sensorimotor cortex
MEP: motor-evoked potential
MI: motor imagery
FDI: first dorsal interosseous
ADM: abductor digiti minimi
SD: standard deviation
rmANOVA: repeated-measures ANOVA
LI: laterality index
ciCOH: corrected imaginary part of coherence
fMRI: functional magnetic resonance imaging
CST: corticospinal tract
EMG: electromyogram
FFT: fast Fourier transform

## Acknowledgements

The authors would like to thank Kohsuke Okada for experimental assistance and to appreciate Sayoko Ishii, Kumi Nanjo, Yoko Mori, Yumiko Kakubari, Shoko Tonomoto, and Aya Kamiya for their technical support during the study.

