## Supplementary files for "Spatially bivariate EEG-neurofeedback can manipulate interhemispheric rebalancing of M1 excitability"

#### Data compliance

In all five sessions, participants subjectively confirmed that they were able to perform kinesthetic MIs of the targeted muscle. EEG signals with amplitudes ≤ 100 μV were consistently recorded throughout the sessions. SMR amplitudes were greater during the resting epoch than during the MI epoch, irrespective of the imagined movement, indicating desynchronization of the sensorimotor neural activity induced by MI. The averaged power spectrum density from the C3 channel during the resting epoch showed peak frequencies at 8–13 Hz and 21–24 Hz across the participants (Crone et al., 1998; Neuper et al., 2006; Pfurtscheller, 2001). The averaged powers in the alpha and beta bands were $1.67 \times 10^{-4} \pm 0.34 \times 10^{-4}$ $V^2$ and $2.85 \times 10^{-5} \pm 0.54 \times 10^{-5}$ $V^2$ (mean ± standard deviation [SD]), respectively. All procedures were well tolerated and no adverse events were noted.

We confirmed the difficulties of each session. The mean value of the sum of the triggered trials (± 1 SD) of all sessions were 14.2 ± 4.7 trials and a post-hoc paired t-test following a rmANOVA revealed no significant difference between the three sessions (all $p > 0.05$; REST: 12.8 ± 4.6 trials, NoFB: 27.7 ± 5.8 trials, HIGH: 12.0 ± 5.0 trials, MID: 16.1 ± 3.8 trials, LOW: 15.9 ± 5.4 trials). The waiting time for a triggered event from MI onset was confirmed for each session. The mean waiting time (± 1 SD) was 1.96 ± 0.81 s, and a post-hoc paired t-test following a rmANOVA showed no significant difference between the sessions (all $p > 0.05$; REST: 1.93 ± 0.82 s, NoFB: 1.22 ± 1.62 s, HIGH: 2.16 ± 0.76 s, MID: 2.05 ± 0.73 s, LOW: 1.71 ± 0.94 s).

#### IHI curves

To validate IHI measurement under bi-EEG-triggered dual-TMS setup, IHI curves were obtained (Supplementary figure 1A) in 20 out of 24 participants, with CS of varying intensity (five different intensities, 100–140% of RMT, in steps of 10% RMT). A one-way rmANOVA for intensities (six levels: 0% [TS only], 100%, 110%, 120%, 130%, and 140% RMT) revealed significant difference in intensity ($F_{(5,95)} = 8.31$, $p < 0.001$, $\eta^2 = 0.28$; 0% RMT [TS only]: 0.93 ± 0.25 mV, 100% RMT: 0.66 ± 0.29 mV, 110% RMT: 0.62 ± 0.31 mV, 120% RMT: 0.54 ± 0.26 mV, 130% RMT: 0.47 ± 0.25 mV, and 140% RMT: 0.43 ± 0.22 mV). A post-hoc two-tailed paired t-test showed significant difference between TS only and 100%–140% of RMT (TS only versus 100% RMT: $p = 0.039$, TS only versus 110%-140% RMT: all $p < 0.001$). There were no significant differences across CS intensities, while the size of MEP amplitude was smaller for larger CS intensities. IHI was approximately half of the maximum when the CS intensity was 120–130% of RMT, which was compatible with a previous EEG-TMS experiment (Stefanou et al., 2018; Tsutsumi et al., 2012). Furthermore, the $SI_{1mV}$ of CS showed approximately 50% of the mean conditioned MEP over the mean unconditioned test MEP (Supplementary figure 1B).

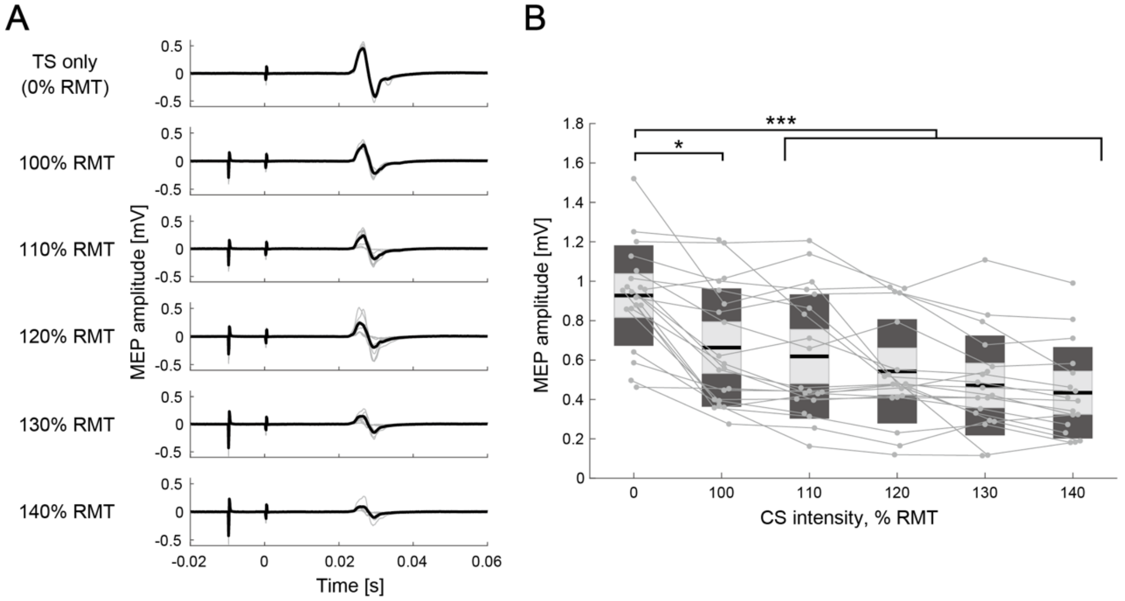


**Supplementary figure 1 IHI curves**

**(A)** MEP amplitudes in the different intensity of conditioning stimulus of a representative participant. The thin gray lines represent each trial and black lines indicate the trial mean. **(B)** The IHI curves of the individual participants are represented by thin gray plots and lines. The figure presents the individual data as an alternative to a box plot. The light grey box represents 1.96 SEM (95% confidence interval) and dark grey box indicates 1 SD. The black line indicates the group mean. The y-axis indicates raw MEP amplitude against CS intensity (x-axis, in %RMT). IHI was approximately half of the maximum when the CS intensity was 120–130% of RMT. Dendrograms above the bars represent the results of the post-hoc analyses. * $p < 0.05$ and *** $p < 0.001$; all comparisons were Bonferroni corrected.

### IHI manipulation for control muscle

In the control muscle (ADM), one-way rmANOVA for sessions (five levels: REST, NoFB, HIGH, MID, and LOW) revealed significant differences ($F_{(4,88)} = 2.51$, $p < 0.048$, $\eta^2 = 0.12$; REST: 74.1 ± 21.5%, NoFB: 94.9 ± 15.7%, HIGH: 84.0 ± 22.6%, MID: 86.0

± 19.4%, LOW: 82.9 ± 22.8%). Across the three BCI-based neurofeedback sessions (i.e., HIGH, MID, and LOW sessions) for the comparison of IHI magnitude, a post-hoc two-tailed paired t-test showed no significant difference between sessions (HIGH-MID: difference = 2.0, Cohen's $d$ = 0.10, $p$ = 1.00; HIGH-LOW: difference = –1.1, Cohen's $d$ = 0.05, $p$ = 1.00; MID-LOW: difference = –3.1, Cohen's $d$ = 0.15, $p$ = 1.00), whereas significant difference was observed between REST and NoFB sessions (difference = 20.8, Cohen's $d$ = 0.45, $p$ = 0.024; Supplementary figure 2A). Statistical analysis for TS-only revealed no significant differences in MEP amplitude between the sessions ($F_{(4,88)} = 0.62$, $p = 0.649$, $\eta^2 = 0.03$; REST: 0.54 ± 0.51 mV, NoFB: 0.50 ± 0.39 mV, HIGH: 0.73 ± 0.61 mV, MID: 0.59 ± 0.50 mV, LOW: 0.65 ± 0.51 mV; Supplementary figure 2B), indicating that IHI manipulation occurred only in the targeted muscle.

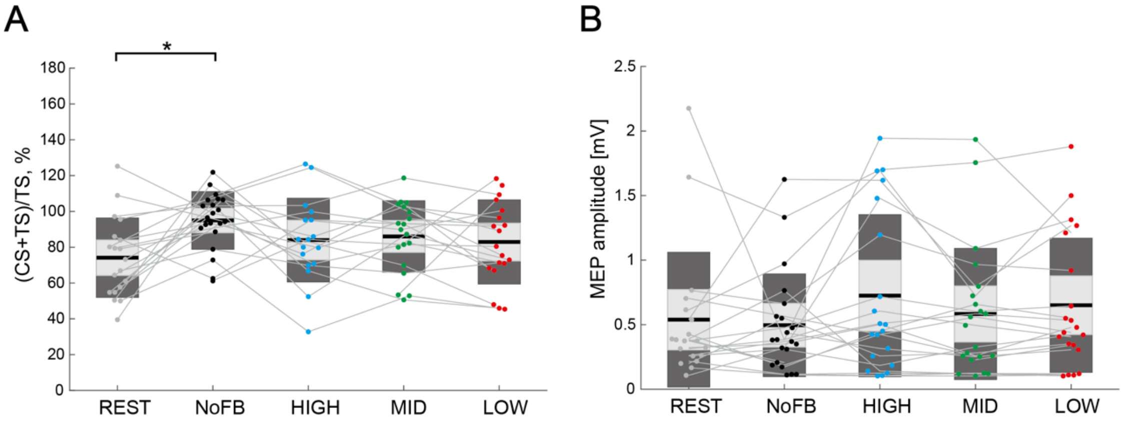


**Supplementary figure 2 Comparison of IHI magnitude for control muscle**

**(A)** The IHI magnitude of the individual participants are represented with colored plots and thin grey lines. The light grey box represents 1.96 SEM (95% confidence interval) and the dark grey box indicates 1 SD. The black line indicates the group mean and colored plots indicates each session. Lower values represent greater inhibitory effect from the ipsilateral hemisphere and therefore more acceleration of the IHI magnitude. Dendrograms above the bars represent the results of the post-hoc analyses. * $p < 0.05$; all comparisons were Bonferroni corrected. **(B)** The figure shows MEP amplitude elicited by a single TS (TS-only). No significant difference in MEP amplitude was observed across the sessions (all $p > 0.05$).

### Comparison of normalized IHI magnitude

To further examine the IHI magnitude across sessions based on individual intrinsic IHI magnitude during MI, we compared the differences between REST, HIGH, MID, and

LOW sessions by normalizing IHI magnitude to the baseline (i.e., NoFB session). A one-way rmANOVA for sessions (four levels: REST, HIGH, MID, and LOW) revealed significant differences ($F_{(3,66)} = 8.32$, $p < 0.001$, $\eta^2 = 0.24$; REST: -15.9 ± 29.8%, HIGH: -27.3 ± 26.5%, MID: 2.7 ± 29.9%, LOW: 17.7 ± 36.2%). Across three BCI-based neurofeedback sessions (i.e., HIGH, MID, and LOW sessions) for the comparison of IHI magnitude, a post-hoc two-tailed paired t-test showed significant differences between HIGH and MID sessions (difference = 30.0, Cohen's $d = 1.06$, $p = 0.017$), and between HIGH and LOW sessions (difference = 45.0, Cohen's $d = 1.42$, $p < 0.001$), but not between MID and LOW sessions (difference = 14.9, Cohen's $d = 0.45$, $p = 0.734$; Supplementary figure 3).

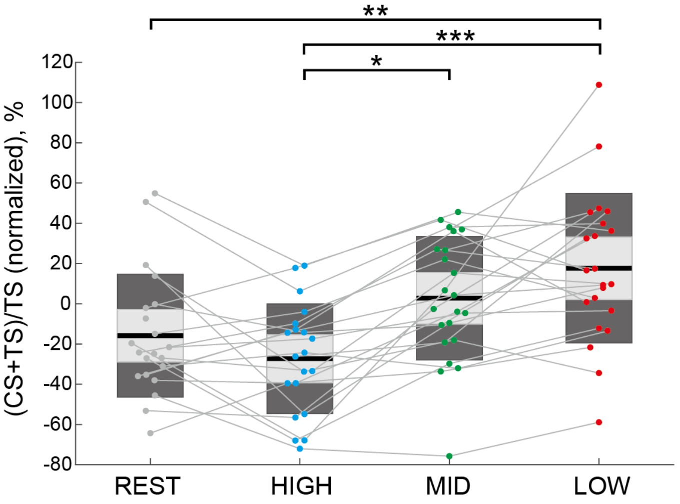


**Supplementary figure 3 Comparison of normalized IHI magnitude**

The normalized IHI magnitude of individual participants are represented by colored plots and thin grey lines. Dendrograms above the bars represent the results of the post-hoc analyses. * $p < 0.05$, ** $p < 0.01$, and *** $p < 0.001$; all comparisons were Bonferroni corrected.

### Comparison of IHI magnitude in non-triggered TMS trials

To examine the influence of spontaneous SMR fluctuations on IHI, non-triggered TMS trial (referred to as the failed trial) was delivered in shuffled timing ranging from 5.5 to 6 s during the MI epoch. For example, data from the HIGH session in Supplementary figure 4 were collected from the MID and LOW sessions, with the target ranges of SMR-ERD in the HIGH session (Supplementary figure 4). A one-way rmANOVA for sessions (three levels: HIGH, MID, and LOW) revealed significant differences ($F_{(2,44)} = 1.43$, $p < 0.248$, $\eta^2 = 0.06$; HIGH: 80.5 ± 34.9%, MID: 83.2 ± 36.3%, LOW: 100.6 ±

46.7%). Although the IHI magnitude in the LOW session was larger than that in the HIGH and LOW sessions (HIGH-MID: difference = 2.7, Cohen's *d* = 0.08; HIGH-LOW: difference = 20.1, Cohen's *d* = 0.49; MID-LOW: difference = 17.4, Cohen's *d* = 0.42;), it was not statistically significant, suggesting that the volitional control of SMR-ERDs in the closed-loop environment differs from spontaneous SMR fluctuations.

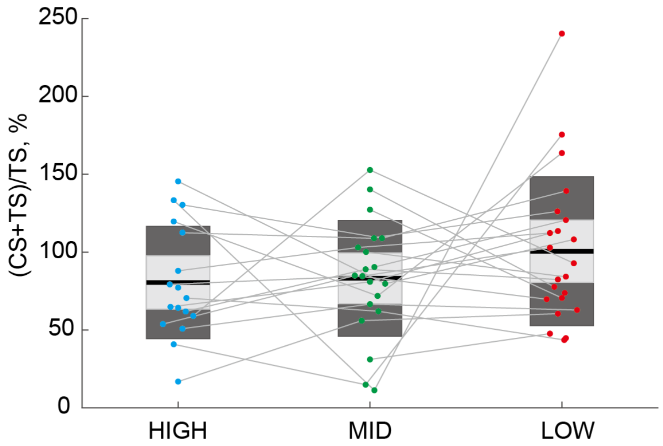


**Supplementary figure 4 Comparison of IHI magnitude in non-triggered TMS trials**

The normalized IHI magnitude of the individual participants are represented by colored plots and thin grey lines. There were no significant differences between sessions in non-triggered trials.

**Connectivity results in the other frequency bands**

In addition to the alpha band, the beta band is also a well-established EEG signature of motor execution and imagery (Crone et al., 1998; Pfurtscheller, 2001). To verify whether resting-state functional connectivity in other frequency bands, i.e., not feedback frequency, was associated with IHI, an across-subject Pearson's correlation was used. In the across-subject correlations between $IHI_{rest}$ and Network-intensity$_{rest}$ in the theta (4-7 Hz), low beta (14-20 Hz), high beta (21-30 Hz), and gamma (31-50 Hz) bands, we found significant correlation in the high beta (r = –0.618, $p$ = 0.004), but not in theta (r = –0303., $p$ = 0.194), low beta (r = –0.229, $p$ = 0.331), and gamma (r = –0.390, $p$ = 0.089; Supplementary figure 5) bands.

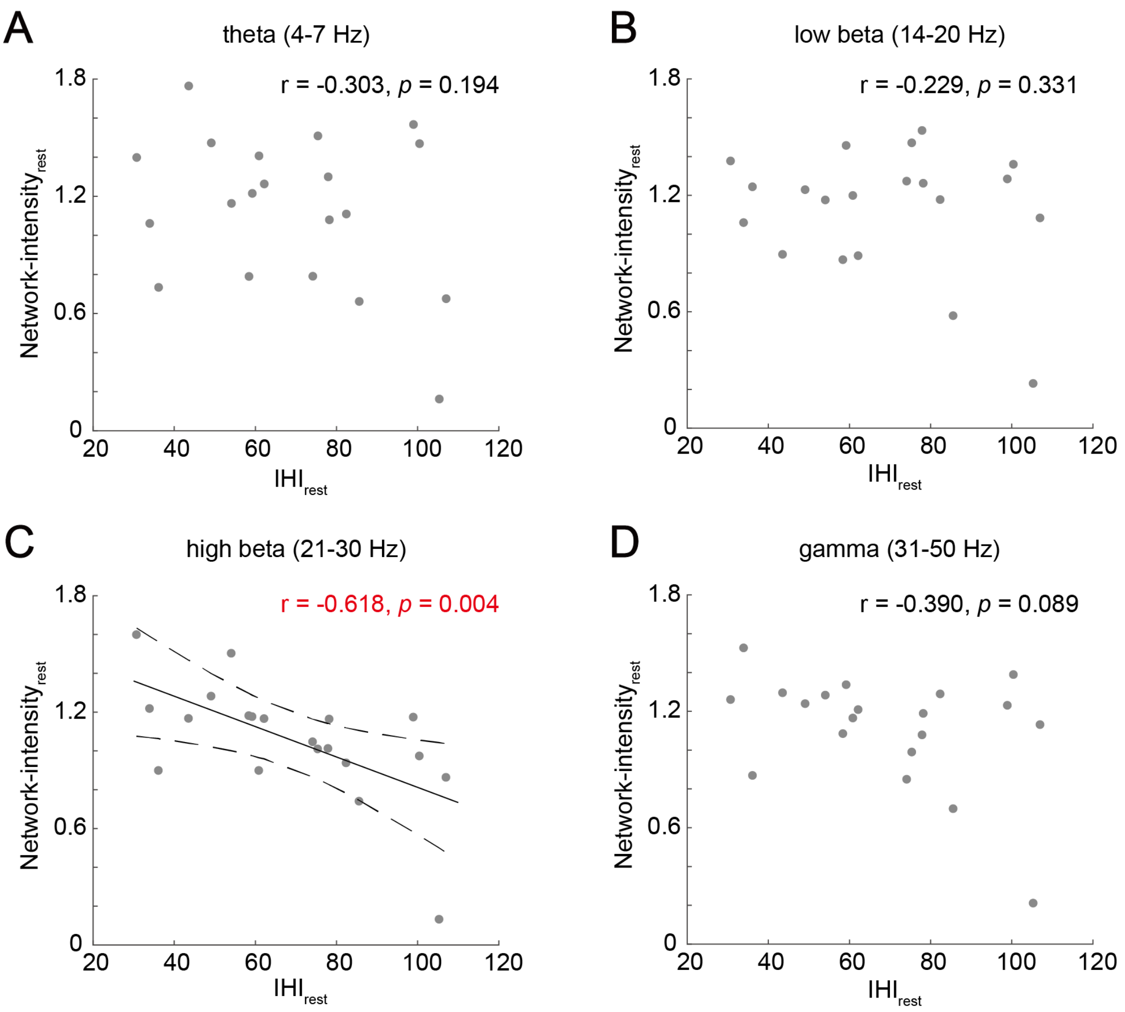


**Supplementary figure 5 Correlation results for the other frequency bands**

Across-subject correlations between IHI at rest and resting-state interhemispheric functional connectivity in the **(A)** theta, **(B)** low beta, **(C)** high beta, and **(D)** gamma bands. Each plot indicates a participant. Solid and dotted lines represent the estimated linear regression and 95% confidence interval, respectively. Only the high beta bands showed significant correlation.
